# Closest relatives of poxviruses replicate in the digestive system of humans and animals worldwide

**DOI:** 10.1101/2024.03.23.586382

**Authors:** Morgan Gaïa, Ulysse Guyet, Hans-Joachim Ruscheweyh, A. Murat Eren, Shinichi Sunagawa, Eugene V. Koonin, Mart Krupovic, Tom O. Delmont

**Affiliations:** Génomique Métabolique, Genoscope, Institut François Jacob, CEA, CNRS, Univ Evry, Université Paris-Saclay, 91057 Evry, France; Research Federation for the study of Global Ocean systems ecology and evolution, FR2022/Tara GOSEE, Paris, France; Department of Biology, Institute of Microbiology and Swiss Institute of Bioinformatics, ETH Zürich, 8093, Zürich, Switzerland; Helmholtz Institute for Functional Marine Biodiversity, 26129 Oldenburg, Germany; Alfred Wegener Institute, Helmholtz Centre for Polar and Marine Research, Bremerhaven, Germany; Institute for Chemistry and Biology of the Marine Environment, University of Oldenburg, Oldenburg, Germany; Marine Biological Laboratory, Woods Hole, MA, USA; Max Planck Institute for Marine Microbiology, Bremen, Germany; Computational Biology Branch, Division of Intramural Research, National Library of Medicine, National Institutes of Health, Bethesda, MD 20894, USA; Institut Pasteur, Université Paris Cité, CNRS UMR6047, Cell Biology and Virology of Archaea Unit,, Paris, France

**Keywords:** Giant viruses, gut microbiome, *Poxviruses*, *Asfuvirales*, eukaryotic viruses, human virus, *Metamonada*, metagenomics

## Abstract

Large and giant double-stranded DNA viruses are diverse and highly prevalent in the environment where they substantially affect the ecology and evolution of eukaryotes^1–4^. Until now, they were only sporadically found in the digestive system of vertebrates^5–7^. Here, we present the discovery of a diversified group of DNA viruses with large linear genomes dubbed egoviruses that comprise a putative viral order most closely related to animal-infecting poxviruses. Egoviruses are found worldwide, almost exclusively in the digestive system of humans, livestock and wild animals. One egovirus is human-specific, evolutionarily constrained, and spread across continents, demonstrating a long-term association between egoviruses and the human population. Analysis of eukaryotic homologs of egovirus core genes suggests that these viruses primarily infect *Trichostomatia* (Ciliates), which play a central role in ruminant digestive processes^8^, and *Parabasalia* (Metamonada), which directly influence animal immunity and gut microbiome ecology^9^. Egoviruses also contributed to the genetic makeup of polintons, endogenous viruses that proliferated in genomes of *Trichomonas* vaginalis^10^ and its close relatives among the *Parabasalia*. Unlike poxviruses that form brick-shaped virions, egoviruses are predicted to form multilayered icosahedral capsids resembling those of asfuviruses and likely representing the ancestral virion structure. Egoviruses currently are the only known clade of eukaryotic DNA viruses abundant in the animal digestive system. Phylogenomic analysis suggests an evolutionary scenario whereby egovirus-like ancestors of poxviruses infected anaerobic unicellular eukaryotes inside the gut of ancestral animals, before transitioning to animal cell infection which was accompanied by the virion metamorphosis.

## Introduction

*Pokkesviricetes* is a class of eukaryotic viruses with large, linear double-stranded DNA genomes within the phylum *Nucleocytoviricota*. The class is divided into two orders, *Chitovirales* (poxviruses) and *Asfuvirales*^11^. Poxviruses infect a broad diversity of animals including humans^12,13^. They include the now eradicated smallpox virus, monkeypox viruses that continue to sporadically infect humans, molluscum contagiosum virus causing benign tumors in immunocompromised individuals, as well as vaccinia virus, a classic model of molecular virology^14–17^. Asfuviruses infect hosts ranging from unicellular eukaryotes to animals^18^ and include the African swine fever virus, which causes deadly and economically important outbreaks in pigs and boars^19^, as well as a wider range of viruses mostly characterized in co-culture with free-living amoeba^20,21^ or directly from the environment (e.g., from plankton) using genome-resolved metagenomics^7,22,23^. Apart from the few viruses of global concern, as declared by the World Health Organization, the class *Pokkesviricetes* is of major importance for our understanding of the diversity, ecology, and evolution of eukaryotic and in particular animal DNA viruses^24,25^.

The evolutionary trajectory of nucleocytoviruses leading to the emergence of animal-infecting poxviruses remains enigmatic. Animals emerged ∼700 million years ago^26^ and host a variety of symbiotic unicellular eukaryotes that for the most part reside in their digestive system^27–30^, representing prime candidates for the ancestral hosts of poxviruses. Indeed, host transition from unicellular eukaryotes to animals would be greatly facilitated by a lasting period of close interactions allowing adaptation to the internal environment of the latter. Yet, decades of investigations using cultivation, PCR surveys, serology, and metagenomics indicated that nucleocytoviruses are mainly transient and thus rarely abundant in the digestive system of humans and other animals^5–7^. Until now, not a single group of *Nucleocytoviricota* was found to be consistently associated with or prevalent in the digestive system, contrasting with the broad distribution and remarkable diversity of these viruses in the environment^4,7,31^. With no evidence of nucleocytoviruses systematically infecting unicellular eukaryotes inside animals, early steps surrounding the emergence of poxviruses could not be explained.

Here, we explored the global diversity and distribution of *Pokkesviricetes* by surveying tens of thousands of metagenomic assemblies from a broad range of environmental samples and animal-associated microbiomes. We discovered a third *Pokkesviricetes* order which we propose naming *Egovirales*. Egoviruses have large genomes and are predicted to form multilayered icosahedral capsids. They are the closest relatives of poxviruses and almost exclusively occur in the digestive system of humans, livestock, and wild animals worldwide. One egovirus population is abundant in ∼1% of the human population across continents. Highly conserved homologs of egovirus core genes were identified in genomes of *Trichostomatia* (Ciliates), *Parabasalia* (Metamonada, including the pathogen *Trichomonas vaginalis*) and *Parabasalia*-associated polintons, another lineage of DNA viruses that integrate into the host genomes^32^, suggestive of a long history of interactions between egoviruses and these animal-associated unicellular eukaryotes. Egoviruses are among the most abundant eukaryotic DNA viruses naturally occurring in vertebrates. They substantially expand the evolutionary scope of *Pokkesviricetes* and provide critical insights into the emergence of poxviruses, pointing to anaerobic unicellular eukaryotes prevalent in the digestive system as an ancestral replicative niche that favored host transition to animal cells.

## Results

### *Egovirales*: a putative *Pokkesviricetes* order characteristic of the digestive system

DNA–dependent RNA polymerases are evolutionarily informative gene markers present in most viruses of the phylum *Nucleocytoviricota*, including all *Pokkesviricetes*^11,33^. We used a comprehensive hidden Markov model (HMM) for the largest RNA polymerase subunit A (RNApolA)^33,34^ to search 85,123 metagenomic assemblies covering a wide range of environmental samples as well as thousands of samples from the digestive systems of humans (e.g., ^35–41^), livestock and wild animals (e.g., ^42–50^) (Table S1). This search identified 1,969,342 RNApolA genes, which we linked to main cellular and viral taxa using a custom reference database (see Methods). Among the *Pokkesviricetes*, we identified a single poxvirus most closely related to *Eptesipox* occurring in the gut microbiome of a marsupial carnivore from Australia^42^ as well as 240 asfuviruses occurring only in the environment, predominantly in aquatic ecosystems (Table S2). These results are in line with the current knowledge of the global niche partitioning of *Pokkesviricetes* and contribute, alongside previous metagenomic surveys^7,23^, to the major expansion of the known diversity of asfuviruses in the environment.

In addition to poxviruses and asfuviruses, we identified a clade of previously undescribed DNA viruses, related to these two viral orders, that were prevalent in the digestive system and which we provisionally name egoviruses. We characterized 12 complete, non-redundant egovirus genomes (average nucleotide identity <98%) from the digestive system of humans, buffalos, goats, chicken, deer, and pigs. These linear genomes are 160-360 kp in length and contain 55 bp inverted terminal repeats, a feature characteristic of *Pokkesviricetes*^51–53^. The egovirus genomes encompass all eight *Nucleocytoviricota* hallmark genes^11,33^ (Table S3), with the family B DNA polymerase coding region often split into two separate open reading frames, echoing trans-splicing of another critical gene (the terminase) recently discovered in mirusviruses^54^. We built an HMM for the egovirus double jelly-roll major capsid protein (DJR-MCP) to complement the RNApolA survey and further expanded the search for egoviruses across metagenomes. Overall, we identified egovirus RNApolA and/or DJR-MCP in 864 metagenomic assemblies, mainly from digestive systems of a wide diversity of vertebrates (Table S3). Using these two marker genes, we recovered more than 200 metagenome-assembled genomes (MAGs) in addition to the 12 complete genomes (Table S3, Supplementary information). Egovirus genomes were on average 81% complete (minimal quality score 50%; see Methods) and were up to 467 kb in size (mean of 229 kb). Phylogenetic analysis of manually curated hallmark proteins confidently identified egoviruses as a distinct major clade within the class *Pokkesviricetes* that is most closely related to poxviruses (Figure 1 and Figure S1). Based on these findings, egoviruses should represent a third order within *Pokkesviricetes* that we propose naming *Egovirales*. *Egovirales* is the only clade within the phylum *Nucleocytoviricota* found to be consistently present in the digestive system.

**Figure 1:**
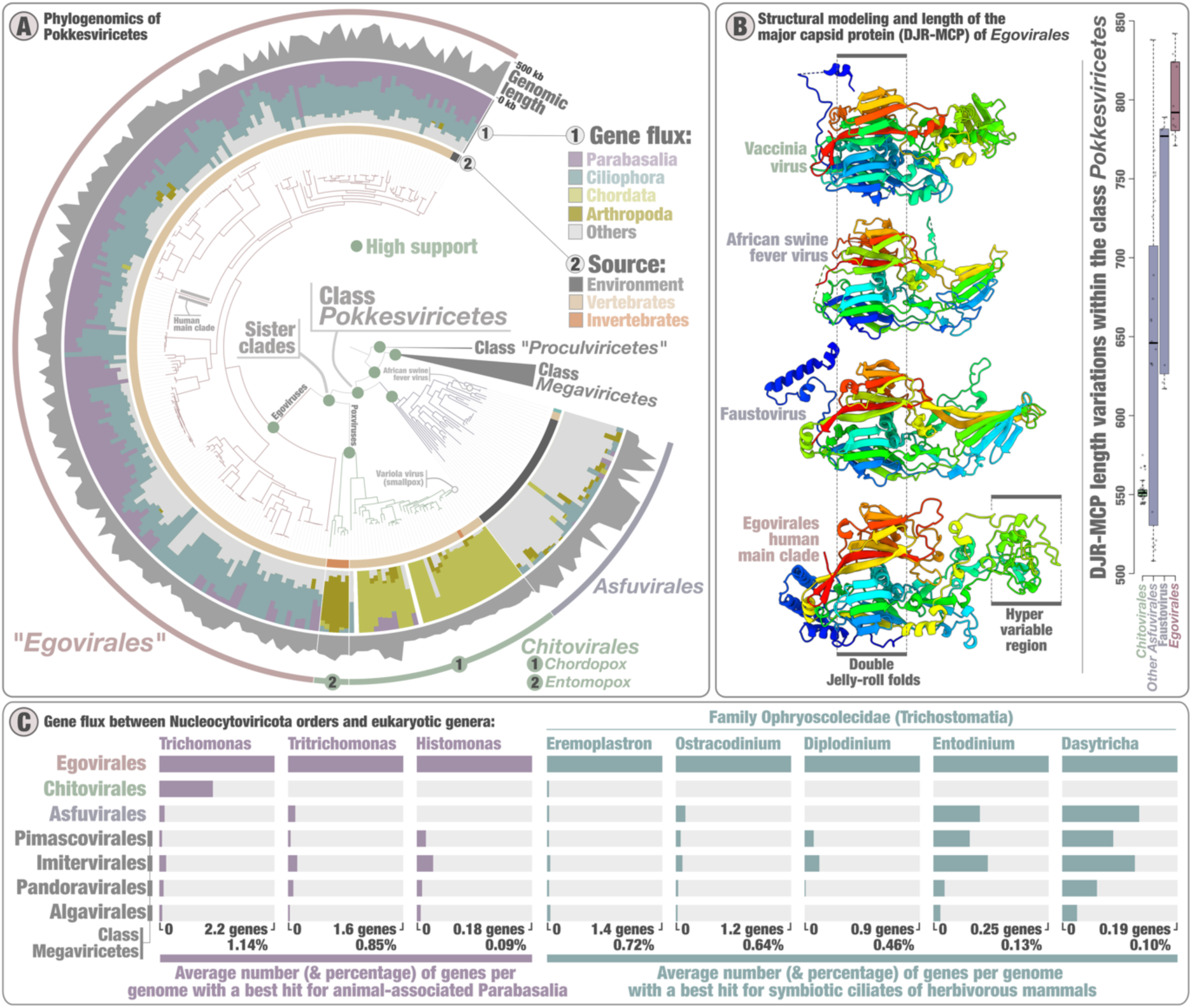
Evolution and predicted gene functions of egoviruses. Panel A displays a maximum-likelihood phylogenomic analysis of members of *Nucleocytoviricota*, along with 202 *Egovirales* genomes with a quality score above 50%. The tree covers 1,807 genomes and was built using concatenated multiple alignments of four hallmark proteins of *Nucleocyovoricota* (DNApolB, RNApolA, RNApolB, TFIIS; 4,110 sites) using IQTree with the LG+F+R10 model. The tree was rooted with the putative class *Proculviricetes* and visualized using anvi’o v8. Green dots in the tree highlight high supports for key nodes (approximate likelihood ratio (aLRT) ≥ 80 and ultrafast bootstrap approximation (UFBoot) ≥ 95; see Methodds). Panel A displays structural modeling of an *Egovirales* DJR-MCP (human main clade) and comparison to the structures of DJR-MCPs from three reference *Pokkesviricetes* DJR-MCPs (from top to bottom: 3sam, 6ku9, 5j7o). The panel also displays boxplots summarizing variation in protein length for the DJR-MCPs of egoviruses (complete genomes), faustoviruses, other clades of asfuviruses, and *Chitovirales*. Panel C summarizes gene flux signal between orders of nucleocytoviruses and eight eukaryotic genera (only the best hits are considered).

We identified 58 core genes represented in at least 10 of the 12 complete *Egovirales* genomes and functionally annotated most of them by sequence analysis and structural modeling (see Methods, Figure S2 and Table S4). These core genes encompass key functionalities of nucleocytoviruses^7,23,34^ and include a histone doublet likely involved in genome condensation into nucleosome-like structures. Histones, including histone doublets, have been previously described in some members of the *Nucleocytoviricota,* in particular, marseilleviruses^55,56^ and medusaviruses^57^, but not in members of *Pokkesviricetes*^58^. Egoviruses encode the major proteins involved in virion morphogenesis that are conserved across *Nucleocytoviricota*, including poxviruses and asfuviruses (Figure S2). The core components of this module are the DJR-MCP^59^ (Figure 1), the A32-like genome packaging ATPase, and the Ulp1-like capsid maturation protease. The *Egovirales* DJR-MCPs are unusually large (average of 806 amino acids among the complete *Egovirales* genomes) and the region facing the exterior of the capsid is highly diversified among egoviruses (Figure S3). These trends echo previous observations on the large MCPs of faustoviruses from the order *Asfuvirales*^20^. In addition, egoviruses encode homologs of the major virion proteins exclusive to asfuviruses, namely, structural polyproteins pp220 and pp62 (see Supplementary information). In African swine fever virus, both polyproteins are processed into multiple structural proteins which play important roles in the formation of multilayered capsids^60,61^. Collectively, the presence of these structural proteins strongly suggests that egoviruses form icosahedral multilayered capsids that are similar to those of *Asfuvirales*^62^, but differ from both the icosahedral capsids of most nucleocytoviruses and the brick-shaped virions of poxviruses^63^.

### Egoviruses are spread worldwide in the digestive systems of vertebrates

We used all the RNApolA and/or DJR-MCP genes of *Egovirales* found in metagenomic assemblies to perform a global survey of the occurrence and diversity of egoviruses (Table S3). Only 9 egovirus-like RNApolA genes were retrieved from environmental samples, including an aquifer^64^, a cold sweep^65^, and marine sediments^66^ (Figure 2). In the phylogenetic tree, these genes formed four small clades, two of which (from marine sediments) were the most basal branches of *Egovirales*. All the remaining RNApolA genes (>97% of the data) originated from the digestive system, representing the bulk of *Egovirales* diversity. The DJR-MCP phylogeny recapitulated these trends, with >99% of the sequences, in this case, originating from the digestive system (Figure S4).

**Figure 2:**
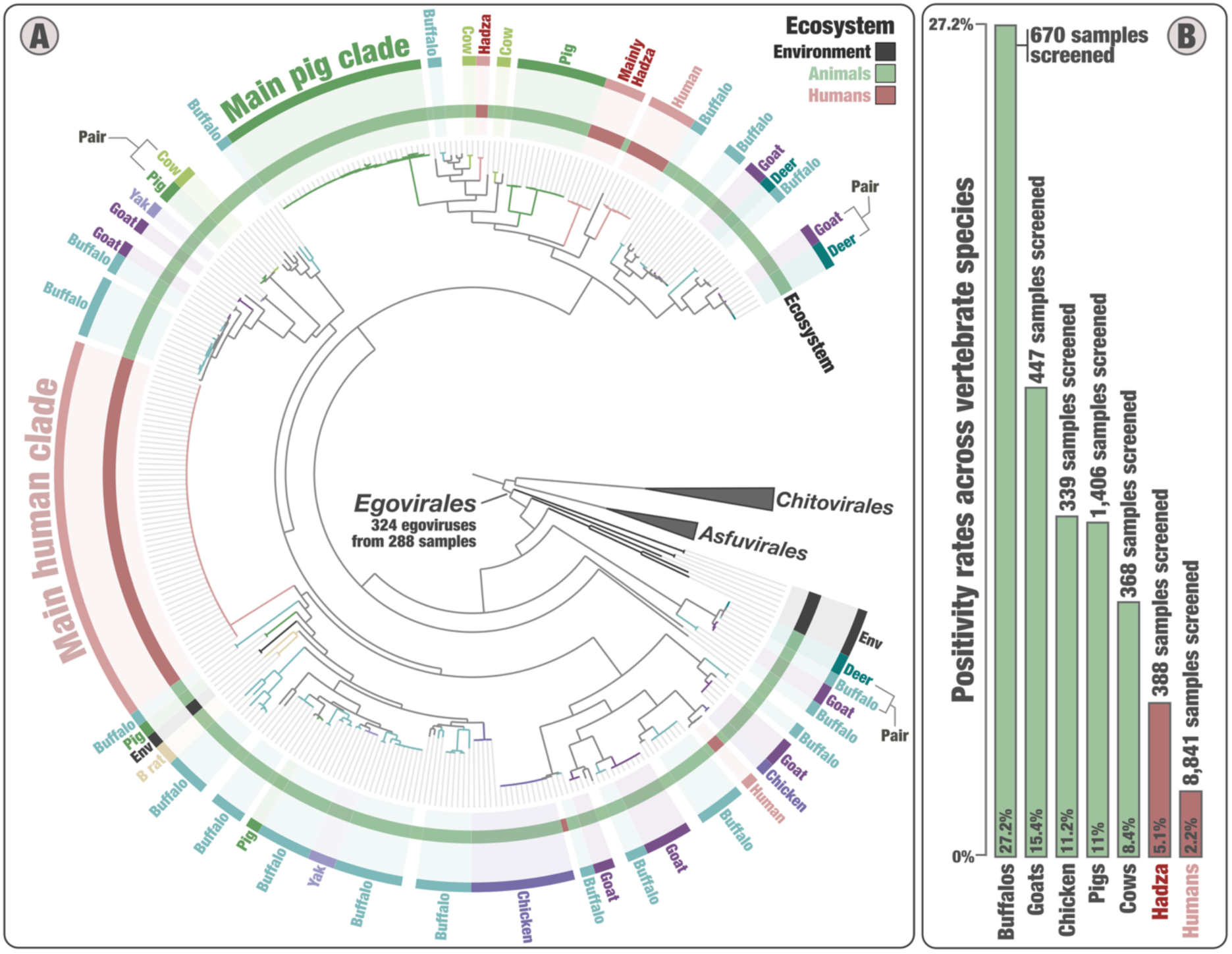
Occurrence of egoviruses among vertebrate species. Left panel displays a maximum-likelihood phylogeny of the RNApolA of egoviruses (n=324), using representatives of *Chitovirales* and *Asfuvirales* for rooting (1,357 sites), and the LG+F+R7 model. Layers of information summarize the occurrence of the egovirus genomes >50 kb in length (including complete genomes), ecosystem type (environment vs. human vs. other animals), and co-occurrence of *Blastocystis* and ciliates. Finally, selections were made for egovirus clades occurring in the same animal species. Right panel summarizes the positivity rate for *Egovirales* (using signal from both the RNApolA and MCP surveys) in cohorts of metagenomic assemblies corresponding to different vertebrate species.

Egoviruses were detected in the digestive system of humans of various age and geographic location (n=217; positivity rate of 2.2% when combining 17 studies encompassing a total of 8,841 human samples; Table S5), with a notable increase of signal among hunter-gatherers from Tanzania (the Hadza; positivity rate of 5.1%) (Figure S5). The extensive sequencing efforts in the study of Hadza populations^35^ could in part explain this enhanced signal. Nevertheless, some of the human-associated clades were either exclusively or mostly detected among the Hadza (Figure 2 and Figure S4), suggesting that lifestyle could affect the diversity of human-associated egoviruses. Egoviruses were also well represented in the digestive system of pigs (n=225; positivity rate 11%), buffalos (n=193; 27%), goats (n=95; 15%), cows (n=44; 8%), chicken (n=38; 11%), and to a lesser extent, also in other livestock for which fewer samples were available (deer, yak, sheep, zebu, reindeer) (Table S3 and Table S5). Buffalos had by far the highest *Egovirales* positivity rate among the surveyed vertebrate species, with viruses covering a wide range of distantly related clades (Figure 2) and occurring abundantly in the ruminant stomach compartments (see Table S6 and Supplementary information). Even fewer metagenomes were available for the digestive system of wild animals, but nevertheless, bamboo rats, a golden bamboo lemur, an olive baboon, and a white rhinoceros were found to be additional carriers of egoviruses, indicating that primates and other wild mammals also represent a substantial reservoir of egovirus diversity.

Examination of the *Egovirales* dispersal indicates that clades of closely related egoviruses tend to occur in the same animal species. This was most apparent for the two main clades associated with humans and pigs (Figure 2 and Figure S4). We characterized the main human clade in 138 metagenomic assemblies from 18 studies that cover various countries from multiple continents, including Tanzania (the Hadza), Nepal, Fiji, USA, England, Sweden, Denmark, Germany, Italy, and Spain (Table S3). Average amino acid identity was 99.4% among the RNApolA genes and 99.9% among the MCP genes, indicating that this clade represents a cohesive viral population occurring among humans at a global scale. A complementary k-mer survey of a MCP gene representative against assemblies of the Sequence Read Archive detected this viral population in 437 metagenomes, broadening the geographic scope of its signal within the human population (Table S5 and Methods). Overall metagenomic signal indicates that this viral population is abundant in more than 1% of the human population (including in remote rural villages and densely populated areas^67,68^), encompasses various stages of the human lifespan (e.g., in a 3-year-old child and a pregnant woman), and remains abundant over time in longitudinal studies of the same individuals (up to nearly one year of sampling). The main pig clade shared similar trends, being found in 109 metagenomic assemblies from 4 studies that cover countries from multiple continents. These results demonstrate that cohesive egovirus populations can cover expansive geographic regions while being strictly associated with a single vertebrate species. In addition, we found instances of closely related egoviruses occurring in multiple vertebrate species. Examples include goat-deer, pig-cow, and deer-buffalo pairs (Figure 2 and Figure S4). Overall, our results demonstrate the worldwide spread and strong co-occurrence of egovirus populations with one or multiple vertebrates species.

### Egoviruses are connected to animal-associated anaerobic unicellular eukaryotes

Repetitive infections over extended periods of time typically result in lateral gene transfers between viruses and their prime eukaryotic hosts, a signal detectable by means of comparative genomics^7,69^. To predict the host range of egoviruses, we performed a global protein-centric genomic comparison between nucleocytoviruses and a wide range of eukaryotic lineages (see Methods). In line with their known host ranges, our results showed a strong gene flux signal between poxviruses and Chordata (chordopoxviruses) or Arthropoda (entomopoxviruses) (Figure 1). In sharp contrast, the same analysis showed strong gene flux signal between egoviruses and two animal-associated groups of unicellular eukaryotes: phylum *Parabasalia* (43% of *Egovirales*-eukaryote gene flux) and order *Trichostomatia* (42%) (Table S7, Supplementary information). This gene flux signal is effectively unique of egoviruses among the known nucleocytoviruses (Figure 1). It is largely driven by egovirus core genes of known function (e.g., family B DNA polymerase, RNA polymerase A, mRNA capping, Holliday junction resolvase) as well as egovirus core genes of unknown function (with no orthologs detected in other nucleocytoviruses), which appear to have been sporadically transferred into *Parabasalia* and/or *Trichostomatia* genomes (see Figures S6-S7-S8 for the corresponding phylogenetic analyses). Within *Parabasalia*, egoviruses were primarily linked to the genera *Trichomonas* and *Tritrichomonas* (Figure 1 and Table S7), which correspond to prevalent pathogens of humans and other animals^70^. Within *Trichostomatia*, the gene flux signal was almost entirely constrained to the family *Ophryoscolecidae* (Figure 1 and Table S7), which is best known for its participation in digestive processes in ruminants (including buffalos)^8^.

Aside from *Parabasalia* and *Trichostomatia*, we did not detect any clear gene flux signal to connect egoviruses with other prevalent animal-associated unicellular eukaryotes (Tables S7-S8-S9). The most notable case in point is *Blastocystis* (a Stramenopile), which is widespread in the digestive system^28,71,72^, coexisting with egoviruses in many human, chicken and pig samples especially within the scope of our metagenomic survey (Figure S5 and Table S5), and yet displays no gene flux to egoviruses. The complete lack of a gene flux signal for *Blastocystis*, despite a considerable niche overlap with egoviruses, quantified here with metagenomics, demonstrates that horizontal gene transfers are not random and do not occur between egoviruses and all co-occurring protists in the digestive system. As a result, the overall gene flux signal and the occurrence of homologs of egovirus core genes in reference *Parabasalia* and *Trichostomatia* genomes (Table S10) appear to be indicative of specific virus-host interactions. Although *Parabasalia* (Metamonada) and *Trichostomatia* (Ciliates) belong to distinct phyla of unicellular eukaryotes^70^, the two groups display commonalities in their lifestyle and cell biology. Namely, both groups of unicellular eukaryotes have anaerobic lifestyle, inhabit the digestive tract of animals and can perform phagocytosis^73,74^, a mechanism well-known to be exploited by large DNA viruses to enter their host cells^75,76^.

### Egoviruses contributed to the genetic makeup of polintons, endogenous viruses of *Parabasalia*

*Parabasalia* genomes, and especially those of *Trichomonas vaginalis*^10^, contain numerous endogenous viral elements, known as polintons, a major lineage of DNA viruses with a DJR-MCP^32^. These endogenous polintons encompass ∼20 genes, one of these encoding KilA, a DNA-binding protein of unknown function that appears to be restricted to polintons integrated in *Parabasalia* genomes^77^. KilA is also a core gene of poxviruses and egoviruses, with a more sporadic occurrence among other nucleocytoviruses of the class *Megaviricetes*. Our phylogenetic analysis of KilA suggests an ancient evolutionary history of this gene among the nucleocytoviruses (Figure 3). This analysis further showed that polinton KilA genes comprised two distinct clades (type I and type II), with their respective common ancestors most likely acquired from distinct egoviruses before the radiation of *Trichomonas*, *Tritrichomonas*, and other related parabasalids from their common ancestor. In the genome of *Trichomonas vaginalis* strain G3 (176 Mb), the two KilA clades are represented by high copy numbers (1,116 copies of type I and 718 copies of type II) and spread across all chromosomes (Figure 3). The second species of *Parabasalia* with the highest copy number of the two clades is that of *Trichomonas tenax*. The apparent transfer of an egovirus core gene to polintons, endogenous viruses of *Parabasalia*, strengthens the connection between egoviruses and this clade of gut-associated anaerobic unicellular eukaryotes.

**Figure 3:**
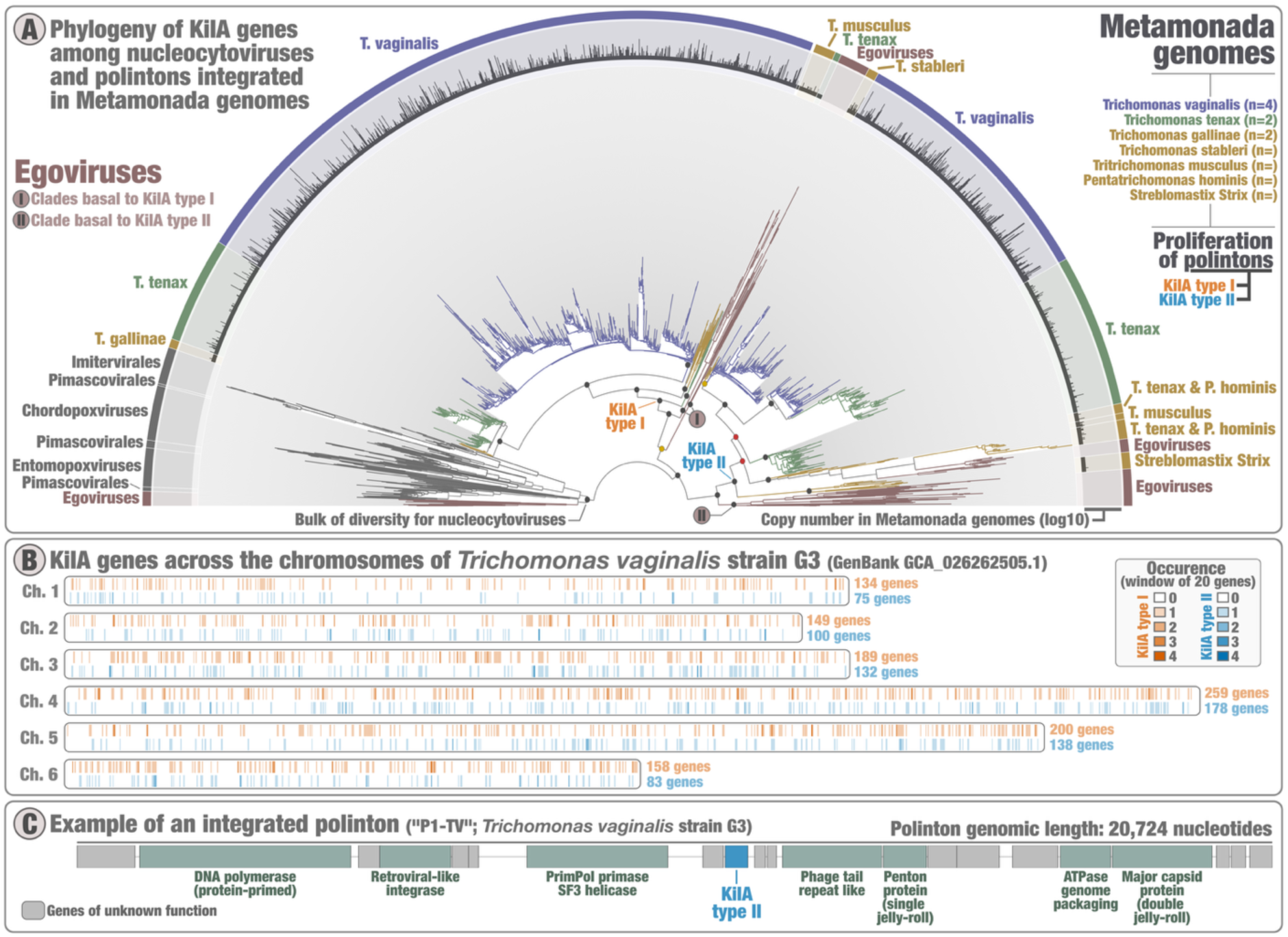
An egovirus origin for the KilA genes of prevalent endogenous polintons in Metamonada genomes. Panel A displays a phylogenetic analysis of KilA predicted proteins from nucleocytoviruses and endogenous polintons (2,207 sequences, 313 sites; model VT+F+R9), which delineates two phylogenetic clades (KilA types I and II) in the polintons. Only representatives of highly redundant Metamonada protein sequences (95% sequence identity) have been selected (see Methods); the number of represented sequences is displayed as “copy number”. Panel B provides the occurrence of the KilA genes across genomes of *Trichomonas vaginalis* strain G3. Panel C displays the genomic structure of polinton P1-TV^77^ integrated into the genome of *Trichomonas vaginalis* strain G3.

## Discussion

Our global metagenomic survey of DNA viruses of class *Pokkesviricetes* led to the discovery of a third putative order in this class, which we provisionally name *Egovirale*s. Egoviruses have linear genomes that can reach 467 kb in length and occur mostly in the digestive system of humans, livestock, and wild animals where they prevail worldwide. Egoviruses have a highly divergent major capsid protein (DJR-MCP) and likely produce viral particles with an icosahedral multilayered structure resembling the virions of African swine fever virus. Egoviruses not only fill major gaps in our knowledge of the ecology and evolution of *Pokkesviricetes*, but also represent the only so far discovered group of viruses with large DNA genomes that is consistently detected in human and animal gut microbiomes. Given its evolutionary trajectory and distribution patterns, this previously overlooked major group of viruses with complex genomes is of prime interest to human and veterinary virology as well as viral evolution.

Until now, the digestive system was considered to encompass a limited diversity and abundance of eukaryotic DNA viruses^5–7^, with the most prevalent ones, such as anelloviruses^78^, having small, single-stranded DNA genomes of only a few kilobases. No convincing evidence of reproduction in the gut is available for any viruses of eukaryotes with large DNA genomes. Egoviruses with their large genomes are changing this view, being projected from the metagenomic assembly signal to be abundant in the digestive systems of ∼180 million humans and billions of livestock and wild animals. These viruses were discovered by metagenome analysis, an approach that only targets the most abundant viral genomes and were detected by this method alone in more than 2% of the analyzed human gut microbiomes and in even higher fractions of many animal microbiomes. The most prevalent human-specific egovirus alone, found in a wide range of metagenomic surveys, is projected to be abundant in the gut of more than 1% of humans. Among animals with highest signal for *Egovirales*, it is also projected that a substantial proportion (possibly, a majority) of the ∼200 million buffalos contain one or multiple egovirus species thriving in their stomach compartments. The diversity of egoviruses is likely to be undersampled in most vertebrate species, including humans. In addition, longitudinal studies indicate that egoviruses can remain abundant over long periods of time in the gut of the same human individual. Thus, egoviruses are diverse, widespread, and resilient viruses prevalent in the digestive system.

Together, comparative genomics and targeted phylogenetic analysis (e.g., of the family B DNA polymerase) depict a strong virus-host interaction between egoviruses and two distantly related groups of unicellular eukaryotes (order *Trichostomatia* and phylum *Parabasalia*) that are both anaerobic, perform phagocytosis, and prevail in the digestive system of vertebrates^29,30,73,74,79,80^. The *Egovirales*-Parabasalia linkage is mostly restricted to *Trichomonas* and *Tritrichomonas*, two genera that include prominent pathogens infecting a wide range of animals and humans and can dramatically impact animal immunity^9^. For instance, *Tritrichomonas foetus* is a parasite of cattle (reproductive tract) and cats (large intestine) as well as a commensal in the stomach and gut of pigs^80^. In addition, *Trichomonas vaginalis* is the most common non-viral sexually transmitted infection among humans with ∼150 million estimated cases per year^81^. The notable case of the apparent egovirus-to-polinton KilA gene transfer followed by the spread of polintons across the genomes of *Trichomonas vaginalis* and *Trichomonas tenax* (parasite inhabiting the human oral cavity^82^) further supports the long-lasting association between egoviruses and *Parabasalia*. The *Egovirales*-*Trichostomatia* linkage is almost entirely restricted to genera of the family *Ophryoscolecidae*, known to play a central role in ruminant digestive processes^8,83^. High prevalence of these *Trichostomatia* and *Parabasalia* genera in vertebrates, and their phagocytosis-centric feeding lifestyle explain the wide spread of egoviruses across vertebrate species and suggest a long history of egovirus replication in the digestive system.

The evolution of poxviruses in insects and vertebrates, including the emergence of smallpox estimated to have killed half a billion humans over the course of a century, has been extensively studied^12,13,17,84^. However, the origin of poxviruses as a group remains enigmatic due to their distant (albeit unequivocal) relationships with other nucleocytoviruses and the lack of intermediate forms. Egoviruses provide that missing link, given their phylogenetic position as the sister clade to poxviruses and their prevalence in the digestive system. The results of this work suggest that the common ancestor of poxviruses and egoviruses infected anaerobic unicellular eukaryotes capable of phagocytosis, most likely, metamonads and ciliates, in the gut of ancestral animals prior to the divergence of insects and vertebrates (protostomes and deuterostomes) from their common ancestor about 700 million years ago^26^. Given that prominent metamonad pathogens, such as *Trichomonas* and *Tritrichomonas*, bind directly to the epithelium during infection^85,86^, and that pathogenic ciliates can also penetrate the animal tissue^87^, it is conceivable that poxviruses have evolved from an egovirus-like ancestor which gained the ability to reproduce in the animal gut epithelial cells. Our analysis of egovirus genomes suggests that the last common ancestor of *Pokkesviricetes* and the common ancestor of egoviruses and poxviruses had a multilayered icosahedral capsid resembling that of asfurviruses. This capsid was replaced by brick-shaped particles in the ancestors of poxviruses following their split from egoviruses and perhaps concomitantly with the invasion of animal gut epithelial cells.

The out-of-the-gut evolutionary scenario for poxviruses seems to align well with the biology of both poxviruses and eukaryotes. On the viral side, entomopoxviruses can perform an early replication phase in the midgut of insects by first infecting the epithelial cells^88^. On the cellular side, phagocytosis is an important component of the immune system of animals^89^ serving as the entry-point for various DNA viruses^75,76^ and is also a major feeding mechanism in both metamonads^90^ and ciliates^91,92^. Thus, the ability of epithelial cells to perform phagocytosis^93,94^ offers one possible mechanistic explanation for how a DNA virus infecting anaerobic unicellular eukaryotes in the digestive system could have successfully initiated the infection of adjacent animal cells. Altogether, this work shows that a distinct group of nucleocytoviruses is common in human and animal gut, in contrast to the previous belief, and sheds light on the likely origin of poxviruses.

## Methods

### Metagenomic database

Metagenomic sequencing datasets were processed as previously described^95,96^ to extract single-copy marker genes from metagenome assembled genomes to build the database for the taxonomic profiling tool mOTUs^97^. Briefly, BBMap^98^ (v.38.71) was used to quality control sequencing reads from all samples by removing adapters from the reads, removing reads that mapped to quality control sequences (PhiX genome) and discarding low-quality reads (trimq=14, maq=20, maxns=1, and minlength=45). Quality-controlled reads were merged using bbmerge.sh with a minimum overlap of 16 bases, resulting in merged, unmerged paired, and single reads. The reads from metagenomic samples were assembled into contigs using the SPAdes assembler^99^ (v3.15.2) in metagenomic mode. Contigs were length-filtered (≥ 500 bp) and gene sequences were predicted using Prodigal^100^ (v2.6.3) with the parameters - c -q -m -p meta.

### A first metagenomic survey using the RNApolA

A broad-spectrum HMM profile for the RNApolA hallmark gene^33^ was searched against the predicted gene sequences from the metagenomic database using hmmsearch (HMMER^101^ 3.3.1) with an e-value cutoff of 1e^-50^. This search resulted in the identification of nearly two million RNApolA genes. Positive hits were then linked back to their source contig.

### Taxonomic affiliation of RNApolA sequences

We collected amino acid sequences corresponding to a broad range of reference RNApolA genes for bacterial, archaeal, eukaryotic, and giant virus lineages characterized by means of cultivation and environmental genomics^34,102,103^. We generated a custom diamond database^104^ (v.2.1.8) from this resource and summarized the taxonomic affiliation of each sequence in a complementary metadata file. We then performed a diamond blast search of RNApolA genes identified from the metagenomic survey using this custom database and exploited the associated metadata file to assess both their taxonomic affiliation and novelty score (based on the percent identity).

### Phylogenetic signal pointing towards egoviruses

We used cdhit^105^ and a 70% identity cut-off at the protein level to remove redundancy among the nearly two million RNApolA genes identified from the metagenomic survey. This step provided a database of 17,293 unique RNApolA proteins longer than 250 amino acids that encapsulate a wide range of cellular and viral diversity recovered from the metagenomic survey. Then, we used the phylogenetic workflow and visualization capabilities of anvi’o^106,107^ (v.8) using muscle^108^ and fast-tree^109^ to identify and remove RNApolA proteins corresponding to bacteria and plastids. Focusing on the 10,853 remaining unique RNApolA proteins, we iteratively used the same anvi’o workflow but this time to identify and remove long branching artifacts. A total of 13 iterations were performed, after which 8,672 unique RNApolA proteins remained. Using this curated database, we explored the phylogenetic signal of RNApolA proteins using taxonomic annotations and the novelty score for guidance. This led to the identification of a monophyletic clade corresponding to the egoviruses, which is most closely related to asfuviruses and poxviruses. We incorporated these sequences in our custom diamond database and performed a second diamond blast on the nearly two million RNApolA proteins, in order to identify *Egovirales* RNApolA proteins (percent identity >50%) and study their source contigs. This methodology recovered 12 complete *Egovirales* genomes.

### Functional annotation of *Egovirales* core genes

We collected reference culture genomes and metagenome-assembled genomes corresponding to *Asfuvirales* and *Chitovirales*, as well as few *Megaviricetes* representatives from the GVDB collection^11^. We added the 13 complete *Egovirales* genomes into this “*Nucleocytoviricota* genomic resource” and used Orthofinder^110^ (V.2.5.5 in “diamond_ultra_sens” mode) to characterize gene clusters. Gene clusters occurring in at least 10 out of the 12 *Egovirales* complete genomes were labelled as “*Egovirales* core genes”, and those occurring as a single copy in at least 10 out of the same 12 genomes were labelled as “*Egovirales* single copy core genes”. Finally, HHpred^111^ profile-profile comparisons and ColabFold^112^ (3D structure predictions and comparisons) were used to functionally annotate the *Egovirales* core genes. In the case of the DJR-MCP of *Egovirales*, ColabFold^112^ successfully predicted its 3D structure and confirmed the presence of the double-jelly roll (DJR) protein components characteristic of the phylum *Nucleocytoviricota*.

### Recovery of eight *Nucleocytoviricota* hallmark genes

We used HMMER^101^ v3.1b2 to search and collect the eight hallmark genes of *Nucleocytoviricota* from the *Nucleocytoviricota* genomic resource (see “*Functional annotation of Egovirales core genes*” section). Given that genomes of faustoviruses from the GVDB collection^11^ had partial DJR-MCP genes, we replaced those with 7 strains of faustoviruses from the NCBI GenBank collection (strains E9, LCD7, M6, S17, liban, vv10, and vv63; https://www.ncbi.nlm.nih.gov/genbank/ accessed February 2024). We also added the Variola virus isolate VARV V563 genome from the GOEV database^34^, and replaced the *Megaviricetes* representatives initially picked from the GVDB collection by others in the GOEV database^34^ based on their distribution as well as the occurrence of hallmark genes. In addition to the RNApolA and DJR-MCP protein sequences already recovered, we obtained sequences for the family-B DNA polymerase (DNApolB), the second largest DNA-dependent RNA polymerase subunits (RNApolB), the transcription factor IIS (TFIIS), the D5-like primase (Primase), the poxvirus late transcription factor (VLTF3) and the packaging ATPase (pATPase). The DNApolB gene, often present on two separate open reading frames in egoviruses, required manual curation through BLASTp alignments (BLAST^113^ v2.10.1) and phylogenetic reconstructions, as previously described^34^. Briefly, when a single egovirus genome had two sequences for this gene, their positions were inspected in a maximum-likelihood single-protein phylogenetic tree, and their sequences were aligned with BLASTp to a close reference sequence and to each other. The two DNApolB sequences for a same egovirus systematically corresponded to a clear split: they were fused and accordingly labelled for further notice. Finally, we created a database of genomes containing at least seven out of the eight hallmark genes, for downstream phylogenetic analyses. For the global phylogenies of the RNApolA and DJR-MCP of egoviruses (Figures 3 and S3), the database was restricted to egoviruses and representatives from the Chitovirales and Asfuvirales for rooting.

### Alignments, trimming and individual gene phylogenetic analyses

From the genomic database created in the previous section (see “*Functional annotation of Egovirales core genes*”), protein sequences corresponding to the same hallmark gene were aligned with MAFFT^114^ v.7.464 and default parameters. The L-INS-i algorithm was used. Sites with more than 70% gaps were systematically trimmed using Goalign v0.3.5 (https://www.github.com/evolbioinfo/goalign). The maximum-likelihood phylogenetic reconstructions were performed using IQ-TREE^115^ v1.6.2, with the ModelFinder^116^ option to determine the best-fitting model. Supports were computed from 1,000 replicates for the Shimodaira-Hasegawa (SH)-like aLRT^117^ and UFBoot^118^. Supports were deemed good when SH-like aLRT ≥80% and UFBoot ≥95%, as per IQ-TREE manual, and moderate when only one of the two conditions was met. The few cases of hallmark genes occurring more than once in each genome were investigated and resolved after removing recent paralogs and clear outliers. All phylogenetic trees (Figure S1 for the eight individual tree phylogenies; Figure 3 and Figure S3 for the global phylogenies of the RNApolA and DJR-MCP of egoviruses) were visualized and rooted using anvi’o.

### A second metagenomic survey using the DJR-MCP

We collected DJR-MCPs from *Egovirales* complete genomes to generate with HMMER^101^ (v3.1b2) an HMM dedicated to the *Egovirales* DJR-MCP. This HMM was then searched against all predicted gene sequences from the metagenomic database, using hmmsearch (HMMER^101^ v.3.3.1) and an e-value cutoff of 1e^-10^. Positive hits were then linked back to their source contig, expanding the scope of *Egovirales*-positive metagenomic samples.

### A database of 224 egovirus MAGs

We analyzed all contigs containing an RNApolA gene or DJR-MCP gene affiliated to *Egovirales* in the context of their associated bins, which were generated using Metabat2^119^ with differential coverage information, as previously described ^95,96^. We removed contigs devoid of any *Egovirales* core gene found to be specific to this order (n=42), a stringent step to only work on high-quality viral MAGs. In addition, we removed MAGs with a quality score (completion minus redundancy) below 50% after creating a collection of 51 *Egovirales* single-copy-core genes and processing the database with anvi’o^106,107^ (v.8.1) to predict gene sequences and functionally annotate hallmark genes using specific HMMs.

### A phylogeny of *Nucleocytoviricota* using concatenated gene alignments

In addition to the single-protein phylogenies, we also performed a multi-protein phylogenetic analysis of the concatenated DNApolB, RNApolA, RNApolB and TFIIS protein sequences. We replaced the few *Megaviricetes* representatives used as outgroup by a much larger genomic database for the non-*Pokkesviricetes* nucleocytoviruses: we used gene markers for the DNApolB, RNApolA, RNApolB and TFIIS corresponding to the classes *Megaviricetes* (exclusion of *Medusavirus* to minimize long-branching artifacts) from the GOEV database^34^, which were all already carefully manually curated. We added to this resource protein sequences for the DNApolB, RNApolA, RNApolB and TFIIS corresponding to all 224 *Egovirales* MAGs. For the DNApolB, RNApolA, RNApolB, and TFIIS, only sequences longer than 100, 200, 200, and 25 amino-acids respectively were kept. This phylogenetic analysis followed the protocol described in the previous section, except that the FFT-NS-i algorithm of MAFFT v7.464 was used for the alignment of each individual protein dataset, before they were trimmed and concatenated. The phylogenetic tree (Figure 1) was visualized and rooted using anvi’o.

### Metagenomic coverage values for egoviruses and co-occurring eukaryotes

We generated a non-redundant database (average nucleotide identity <95%) of egovirus MCP genes, egovirus RNApolA genes, and eukaryotic RNApolA genes characterized from the *Egovirales*-positive samples. We performed read recruitment of all the *Egovirales*-positive metagenomic samples against the database. For that, quality controlled reads were mapped (Bowtie2^120^, v2.4.116) against the database and the ‘anvi-profile-blitz‘^106,107^ routine used alignments to generate mean coverage values. Taxonomic information was included using the diamond blast for RNApolA genes (see “Taxonomic affiliation of RNApolA sequences” section) and the phylogenetic tree for MCP genes (see “Alignments, trimming and individual gene phylogenetic analyses” section).

### K-mer survey of the most prevalent human-related egovirus population against SRA

A global k-mer survey of a MCP gene representative (MAG: ZEEV15_1_SAMEA3664761_BIN_00000011) against the Sequence Read Archive (SRA) data (at the assembly level) was performed using the online tool Logan-Search^121^, with a minimum k-mer coverage of 50%. Because the tool only allows screening of nucleotide sequences smaller than 1 kb, we only analyzed the first 1 kb of the MCP gene representative.

### Protein sequence comparisons between nucleocytoviruses and cellular organisms

Egovirales proteins, together with *Pokkesviricetes* (*Asfuvirales* and *Chitovirales*) from the GVDB collection and *Megaviricetes* from the GOEV collection, were used as queries for similarity searches against UniProt-TrEMBL (UniProt Knowledgebase Release 2024-03^122^) using MMseqs2^123^ with default settings. Hits not clearly assigned to cellular organisms were removed based on taxonomic annotations (see Supp Table S9). For each query, only the best hit with >= 30% identity and >= 100 bit scores was retained. Given the enrichment in Metamonada and Trichostomatia (Ciliophora) among hits for *Egovirales* proteins, the same queries were searched against additional metamonad and ciliophoran genomes retrieved from NCBI Genome and the Genome Warehouse (list in Supp Table S9) using identical parameters. Bit scores were used to compare the hits across the independent runs, and only the highest-scoring hit per query was retained. For functional overview, Egovirales proteins and their corresponding best hits in metamonads and ciliophora were queried for searches against Pfam-A.full (release 38.0) using MMseqs2 (default settings), retaining only the best hit per query. In parallel, the entire set of egovirales proteins was clustered into orthogroups using OrthoFinder (v2.5.5, default settings), and orthogroup assignments were mapped to those from the functional annotation (performed on complete egovirales genomes only). Similarity searches were performed for the opposite direction, using metamonad and ciliophoran sequences matching egovirales proteins (including *Trichomonas vaginalis* sequences from UniProt-TrEMBL) as queries against *Egovirales*, *Pokkesviricetes* (*Asfuvirales* and *Chitovirales*), and *Megaviricetes* protein datasets (MMseqs2, default settings) (Supp Table S10). These sequences were also searched against UniProt-TrEMBL to identify and exclude potential false positives. For this latter, the same taxonomic filtering was applied to remove hits without clear cellular assignation, and hits to *Metamonada* were removed when using metamonad queries to remove self-matches. For each query, only the best hit per target category was retained.

### Phylogenies for egovirus core genes found in the genomes of Parabasalia and Trichostomatia

Our Protein sequence comparisons identified a gene flux signal connecting egoviruses and two clades of eukaryotes: Parabasalia and Trichostomatia. Most of this signal was due to egovirus core genes occurring in the eukaryotic genomes. To clarify the directionality of this gene flux, we performed phylogenetic analyses for four egovirus core genes of known function (DNA-dependent RNA polymerase largest subunit (RNAPa), family B DNA polymerase (DNApolB), mRNA capping enzyme and Holliday junction resolvase) and two egovirus core genes of unknown function (OG0000040 and OG0000089). HMM profiles for the mRNA capping enzyme (PF03919.22) and Holliday junction resolvase (PF04848) were retrieved from the InterPro database (ebi.ac.uk/interpro/), while new HMMs were constructed from aligned egovirales sequences for OG0000040 and OG0000089 using HMMER. Homologous sequences were identified using HMMER and collected from *Pokkesviricetes*, *Megaviricetes*, and the set of additional *Metamonada* and *Ciliophora* (see previous section), as well as from the TCS database (derived from EukProt v3 and representing the diversity of eukaryotes^124^). To reduce redundancy for the mRNA capping enzyme and Holliday junction resolvase, sequences were clustered with CD-HIT with dataset-specific thresholds (>=80 and 95% identity for TCS sequences in the mRNA capping enzyme and Holliday junction resolvase datasets, respectively; >=90% identity for GOEV sequences in both gene families). Sequences substantially shorter than the dataset average were excluded. For DNApolB and RNAPa, HMM profiles and cellular/viral homologs were obtained from a previous study^34^, and homologs from metamonad, ciliophora, and egoviruses identified using HMMER were directly incorporated into these datasets. Phylogenetic trees were reconstructed as described above, and long-branches sequences, ambiguous taxa, and clades highly divergent from egoviruses were iteratively removed prior to final inference in order to improve the resolution and stability.

### KilA linking egoviruses to metamonad polintons

The KilA gene, which is associated with polinton-like elements in metamonads, especially parabasalids, was analyzed separately due to its distinct evolutionary context. An HMM profile (PF04383.19) was retrieved from InterPro and used to identify homologous sequences across Pokkesviricetes (including egoviruses), Megaviricetes, Ciliophora, and Metamonada. Considering the specific enrichment of polinton-like elements in parabasalids, additional genomes were included: *T. vaginalis* strains ATCC 30188 (GCA_002891335.1) and ATCC 50138 (GCA_052324655.1), *T. tenax* isolate Hs4:NIH (GCA_047301845.1) and strains NIH4 ATCC 30207 (GCA_900231805.1), *T. gallinae* isolate XT1081-07 (GCA_008369845.1), T. sp genotypes 1c (GCA_047372395.1) and 2a (GCA_047372375.1), and *Pentatrichomonas hominis* isolate Hs-3:NIH (GCA_047301545.1). To limit redundancy, sequences from GOEV, GVDB, and each ciliophoran and metamonad genomes were clustered independently using CD-HIT (>=95% identity threshold), and only the representative sequence of each cluster was retained for phylogenetic inferences, along with the number of sequences it represents. Gene trees were reconstructed using the same approach as previously described, and long branches and/or ambiguous taxa were iteratively removed.

### Genomic synteny

Putative genes for the genome of *Trichomonas vaginalis* strain G3 (GenBank reference GCA-026262505.1) were retrieved, including their functional annotation and predicted positions of polinton-like regions (maverick, REF doi.org/10.1038/s41467-025-61483-w). Genomic positions of egovirales-like hits as well as KilA genes of type I and type II were mapped onto it. Finally, MCP genes matching that of Polinton-1_TV, a well annotated polintovirus from the same *Trichomonas vaginalis* strain G3^32^ collected from the REPBASE database girinst.org/repbase/), were identified using mmseqs2 and default settings; the hits were marked in the same table.

## Data availability

Databases our study used or generated include (1) the reference RNApolA proteins (with taxonomy) and those identified from the metagenomic survey, (2) *Egovirales* RNApolA and DJR-MCPs proteins, (3) reference *Nucleocytoviricota* HMMs as well as two new HMMs targeting the DJR-MCP and VLTF3 of *Egovirales*, (4) HMMs for the core genes and single copy core genes of *Egovirales*, (5) the 224 *Egovirales* MAGs (including complete genomes), (6) the 3D structure of the *Egovirales* DJR-MCP, (7) the non-redundant database of MCP and RNApolA genes covering the diversity of egoviruses and eukaryotes found in the *Egovirales*-positive samples, (8) KilA proteins found in nucleocytoviruses and Metamonada, (9) and data associated with the main phylogenies. These items along with the supplementary tables and Supplementary Information document have been made publicly available at https://doi.org/10.6084/m9.figshare.25398895.

## Acknowledgments

Our survey was made possible thanks to the large amount of metagenomic data that has been produced and made publicly available in recent years. As a result, we are grateful to all the scientists involved in the metagenomic exploration of a wide range of environmental ecosystems as well as the human and animal gut microbiomes. We are also grateful to Lucas Paoli for helping facilitate the acquisition of results from the initial RNApolA survey as part parallel metagenomic investigations, as well as to Natalya Yutin for having dedicated time to exploring the egovirus-eukaryotic associations from a comparative genomic perspective. This study was supported by two grants from the French national research agency (ANR-23-CE02-0022 and ANR-23-CE02-0025).

## Additional information

Correspondence should be addressed to Tom O. Delmont.

## Supplemental Tables

**Table S1:** Metadata associated with the metagenomic survey.

**Table S2:** Metagenomic signal for *Chitovirales* and *Asfuvirales* based on the RNApolA survey.

**Table S3:** Main statistics for egoviruses. The table includes metagenomic signal for *Egovirales* based on the RNApolA and MCP surveys, as well as main statistics regarding *Egovirales*-positive samples, and egovirus genomics.

**Table S4:** Gene clusters, core genes and functional annotations for the 12 complete *Egovirales* genomes.

**Table S5:** Occurrence and positivity rate of *Egovirales*, Ciliates, and *Blastocystis* among humans, buffalos, chicken, pigs, and cows.

**Table S6:** Metagenomic signal for *Egovirales* in buffalo samples. The survey covers seven Chinese provinces, three buffalo breeds and different segments of their digestive system.

**Table S7: Gene flux signal between nucleocytoviruses (query) and eukaryotes (targets).** The table summarizes gene flux signal (including functional annotations) observed between nucleocytoviruses, mirusviruses and a wide range of eukaryotic lineages, with focused analyses on the linkage with Metamonada and Ciliates. The table also provides gene flux signal at the level of egovirus orthogroups.

**Table S8:** Metagenomic signal for eukaryotes among the *Egovirales*-positive samples based on the RNApolA survey. The table summarizes the assigned taxonomy of eukaryotic RNApolA hits in the context of the egovirus-positive samples.

**Table S9:** Relative abundance of egoviruses and eukaryotes among the *Egovirales*-positive samples. The table summarizes metagenomic coverage values for MCP and RNApolA genes corresponding to egoviruses and eukaryotes found in the *Egovirales*-positive.

**Table S10: Gene flux signal between eukaryotes (query) and nucleocytoviruses (targets).** The table summarizes gene flux signal (including functional annotations) observed between eukaryotes (Metamonada and Ciliates) and nucleocytoviruses. The table also provides gene flux signal at the level of egovirus orthogroups.

**Table S11: Genomic positions and evolutionary context of the gene KilA in Metamonada-associated polintons.** The table summarizes the list of genes KilA found in Metamonada-associated polintons, and the occurrence of KilA genes of type I and type II across the chromosomes of *Trichomonas vaginalis* strain G3.

## Supplemental Figures

**Figure S1:**
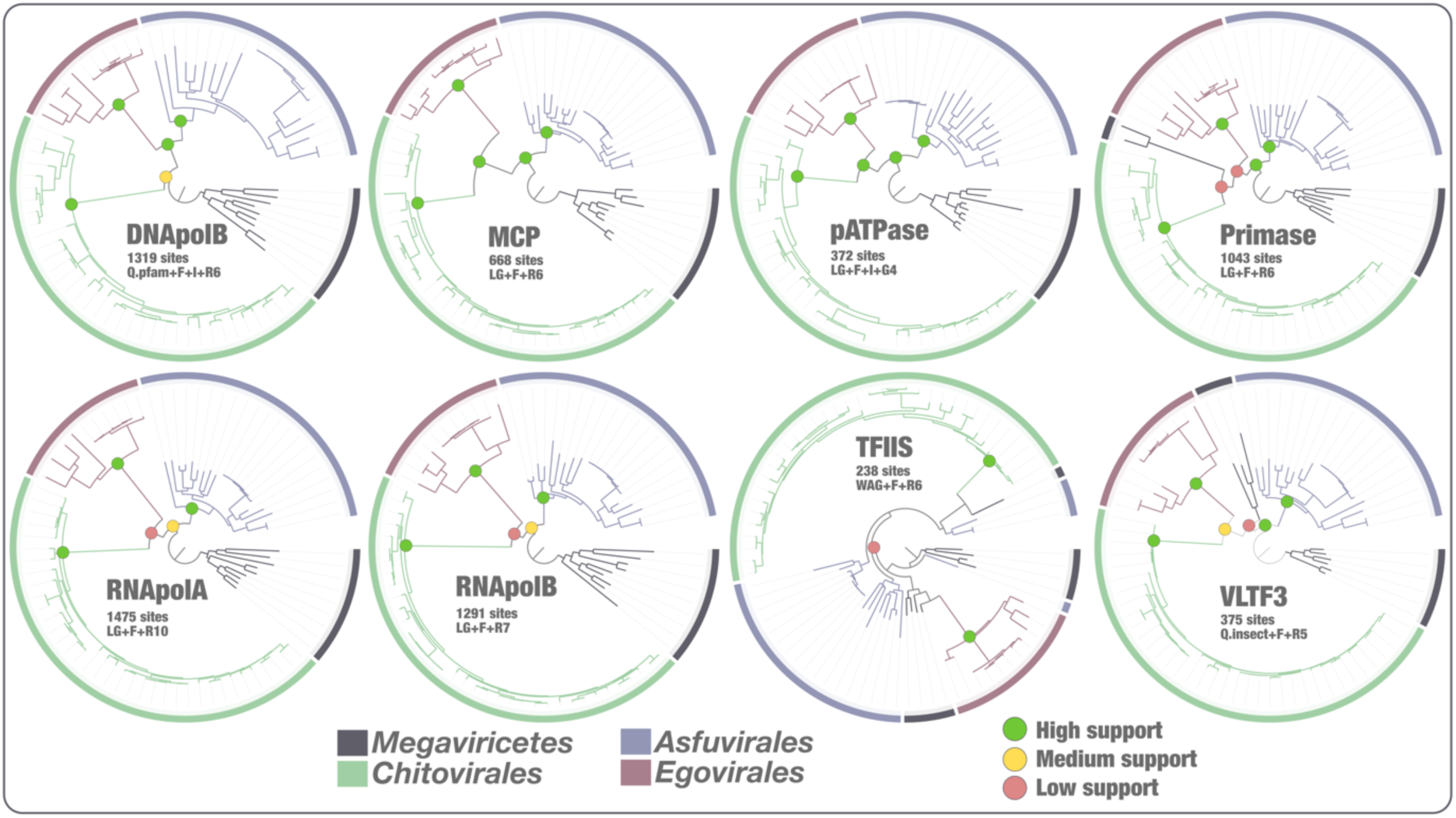
Phylogeny of *Pokkesviricetes* using seven individual hallmark genes. Each tree corresponds to a phylogeny of the same genomic resource (11 *Egovirales*, 36 *Chitovirales*, 21 *Asfuvirales*, and 9 *Megaviricetes*) using one of the eight hallmark genes shared by the tree orders of the class *Pokkesviricetes*. Each genome in this restrained genomic resource contains at least seven out of the eight hallmark genes. Green dots at nodes in the tree highlight high supports (approximate likelihood ratio (aLRT) ≥ 80 and ultrafast bootstrap approximation (UFBoot) ≥ 95; see Methods). Yellow dots represent moderate supports (aLRT >= 80 or UFBoot >= 95).

**Figure S2:**
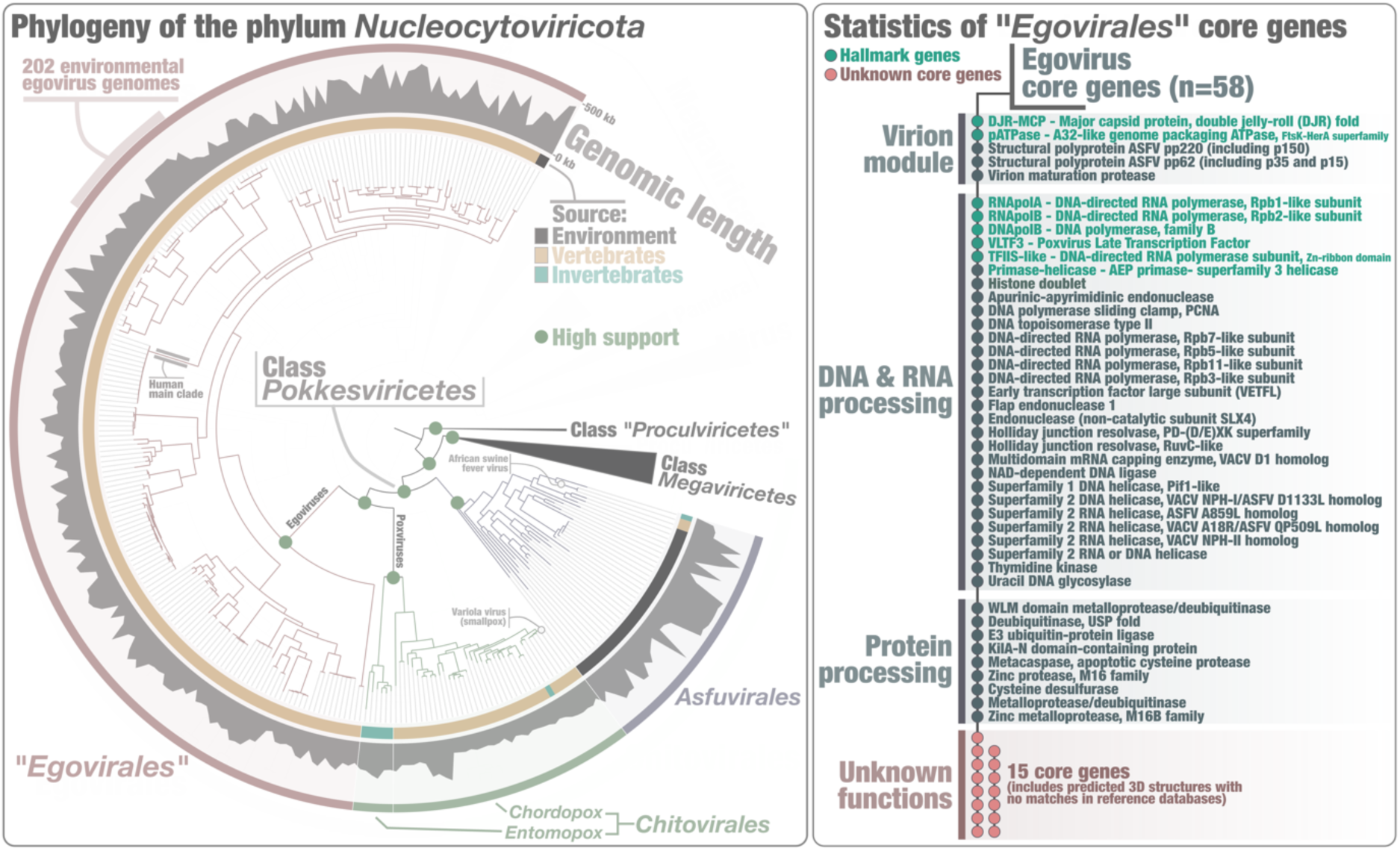
Evolution and predicted gene functions of egoviruses. Left panel displays a maximum-likelihood phylogenomic analysis of members of *Nucleocytoviricota*, along with 202 *Egovirales* genomes with a quality score above 50%. The tree covers 1,807 genomes and was built using concatenated multiple alignments of four hallmark proteins of *Nucleocyovoricota* (DNApolB, RNApolA, RNApolB, TFIIS; 4,110 sites) using IQTree with the LG+F+R10 model. The tree was rooted with the putative class *Proculviricetes* and visualized using anvi’o v8. Green dots at nodes in the tree highlight high supports (approximate likelihood ratio (aLRT) ≥ 80 and ultrafast bootstrap approximation (UFBoot) ≥ 95; see Methods). Right panel summarizes the functional annotation of 58 *Egovirales* core genes.

**Figure S3:**
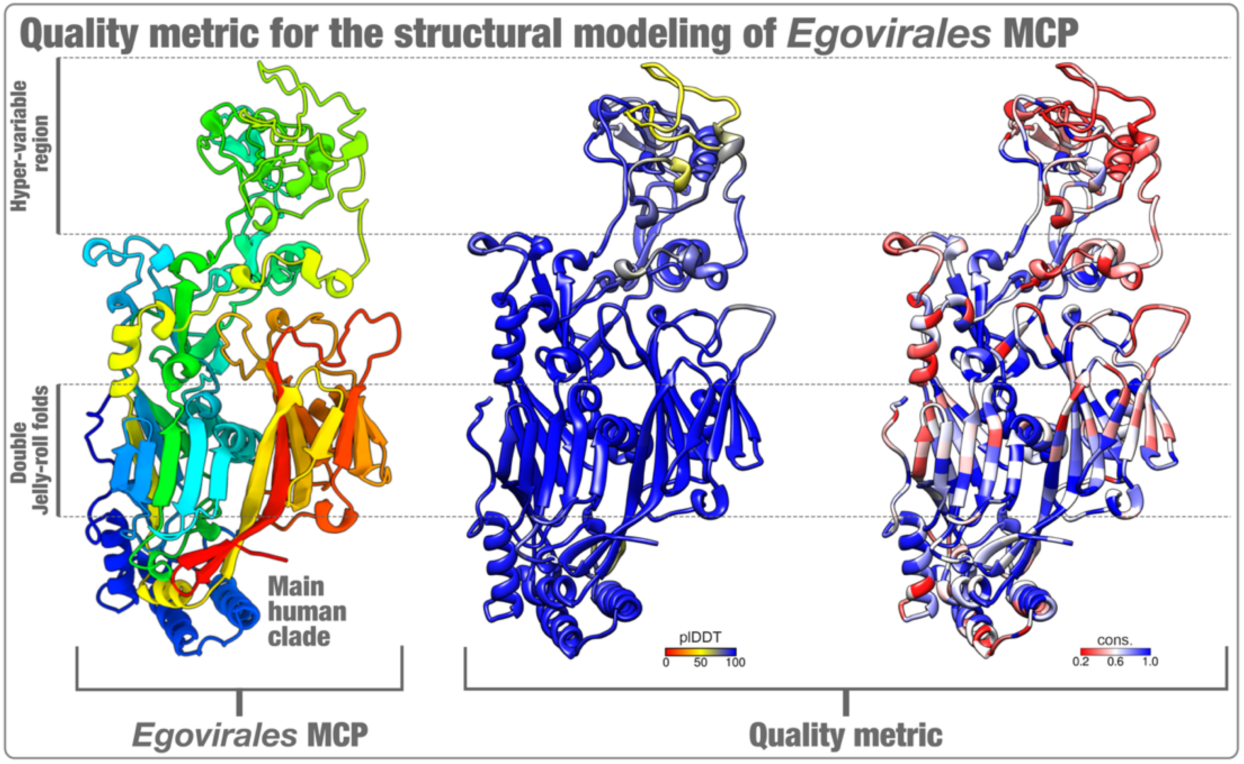
Quality metric of the structural modeling of the Egovirales major capsid protein. Left panel displays the structural modeling of an *Egovirales* DJR MCP (human main clade), as predicted from Colabfold^112^. Right panels display the pIDDT and conservation levels (quality metric).

**Figure S4:**
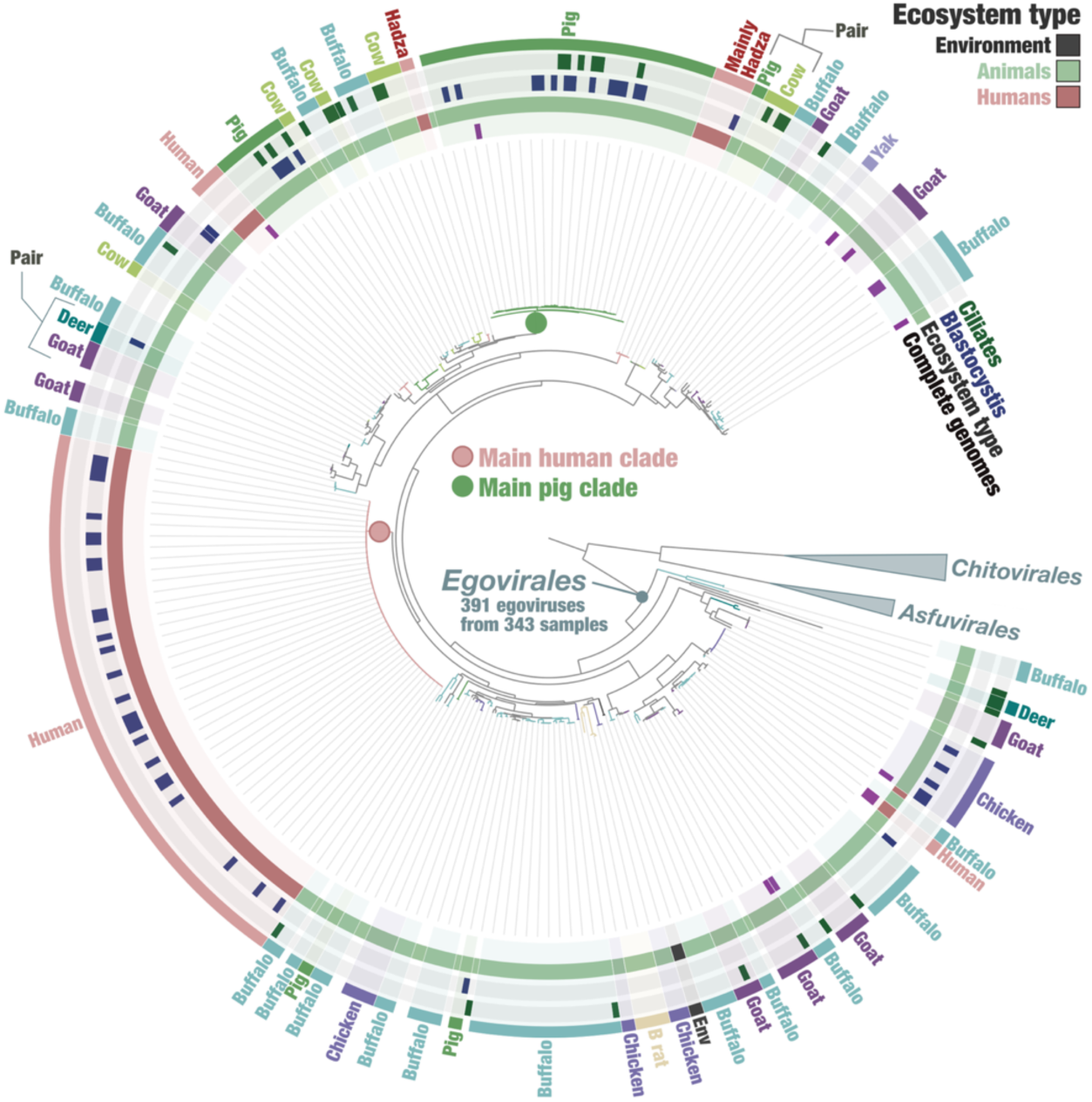
Overlay of the evolutionary tree of egoviruses and their occurrence among vertebrate species. The figure displays a maximum-likelihood phylogeny of the MCP of egoviruses (>600 amino acids; n=391), using representatives of *Chitovirales* and *Asfuvirales* for rooting (784 sites), and the LG+F+R10 model. Layers of information summarize the occurrence of the 12 complete egovirus genomes, ecosystem type (environment vs. human vs. other animals), and co-occurrence of *Blastocystis* and ciliates. Finally, selections were made for egovirus clades occurring in the same animal species.

**Figure S5:**
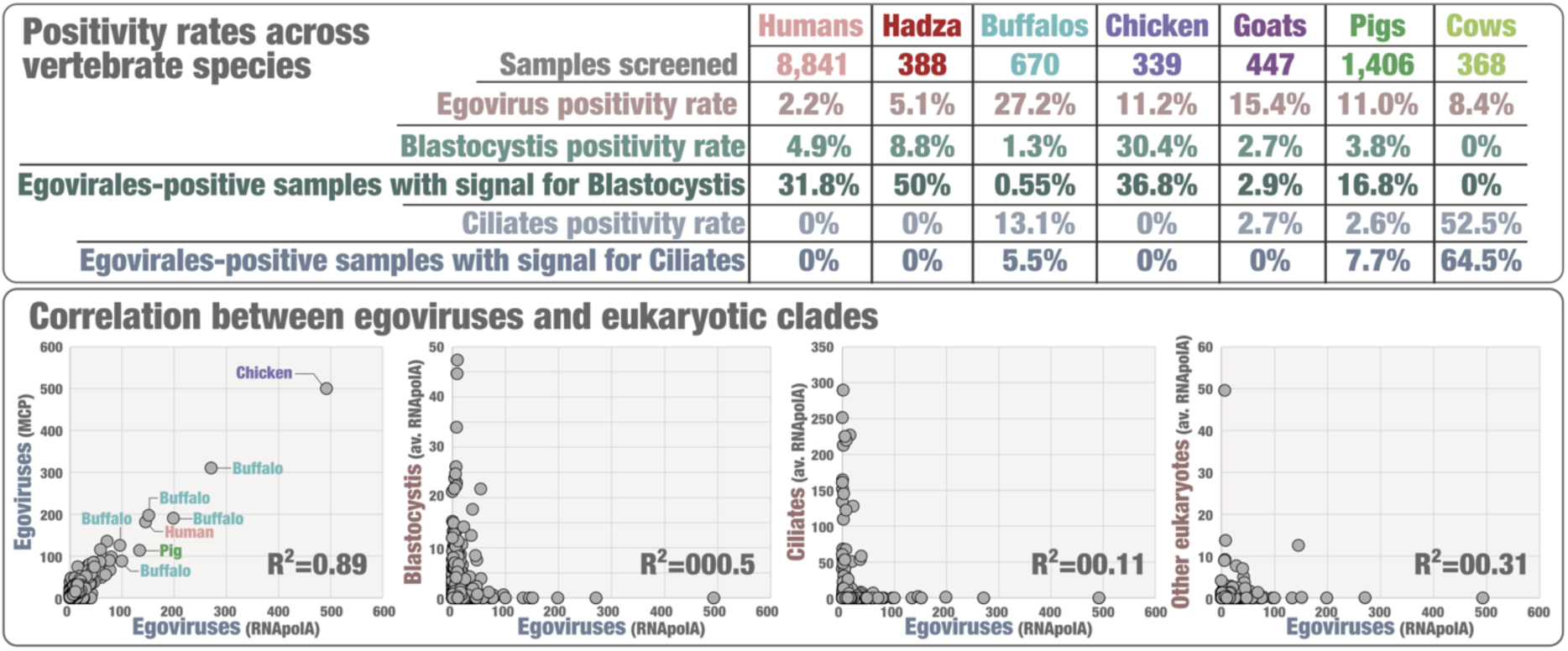
Prevalence of egoviruses and co-occurring unicellular eukaryotes. Top panel summarizes the positivity rate for egoviruses, *Blastocystis* and ciliates in gut metagenomes from five vertebrate species. Bottom panel displays the mean coverage correlation between egoviruses and clades of unicellular eukaryotes across metagenomes corresponding to the 864 *Egovirales*-positive samples.

**Figure S6:**
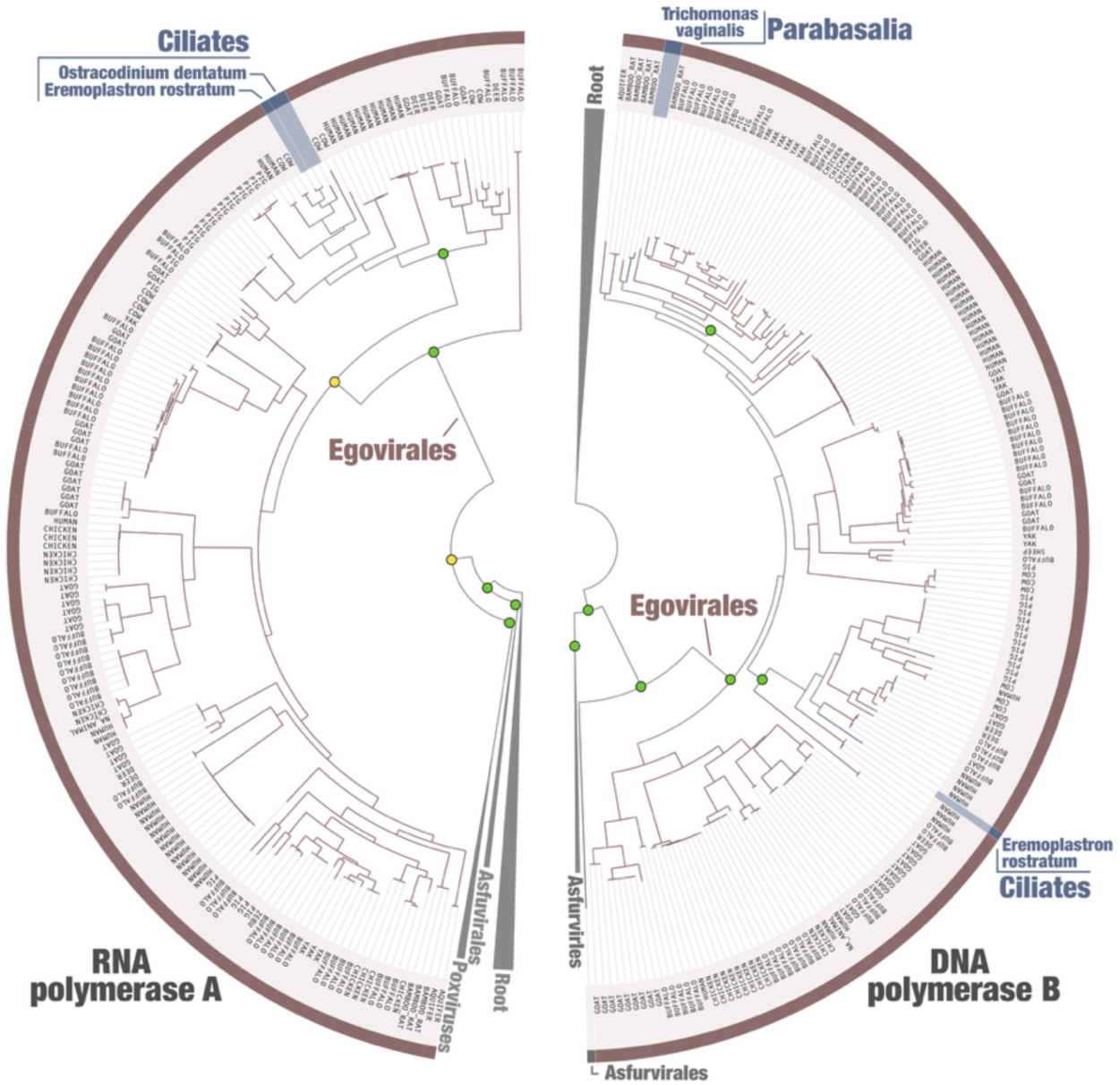
Phylogenies of the RNA polymerase A subunit and DNA polymerase B family. Left tree corresponds to the maximum-likelihood phylogeny of the RNA polymerase A subunit of *Nucleocytoviricota* and eukaryotes (minimum protein length: 200 aa; 1738 taxa; 1,513 sites; model LG+F+R10), rooted between *Pokkesviricetes* (including Egovirales) and the rest. Right tree corresponds to the maximum-likelihood phylogeny of the DNA polymerase B of *Nucleocytoviricota* and eukaryotes (minimum and maximum protein lengths: 100 and 2,000 aa; 485 taxa; 1,255 sites; model Q.PFAM+F+R10), rooted at a level including all sequences except those forming two sister clades for Asfuvirales and egoviruses. Green dots at nodes in the tree highlight high supports (approximate likelihood ratio (aLRT) ≥ 80 and ultrafast bootstrap approximation (UFBoot) ≥ 95; see Methods). Yellow dots represent moderate supports (aLRT >= 80 or UFBoot >= 95).

**Figure S7:**
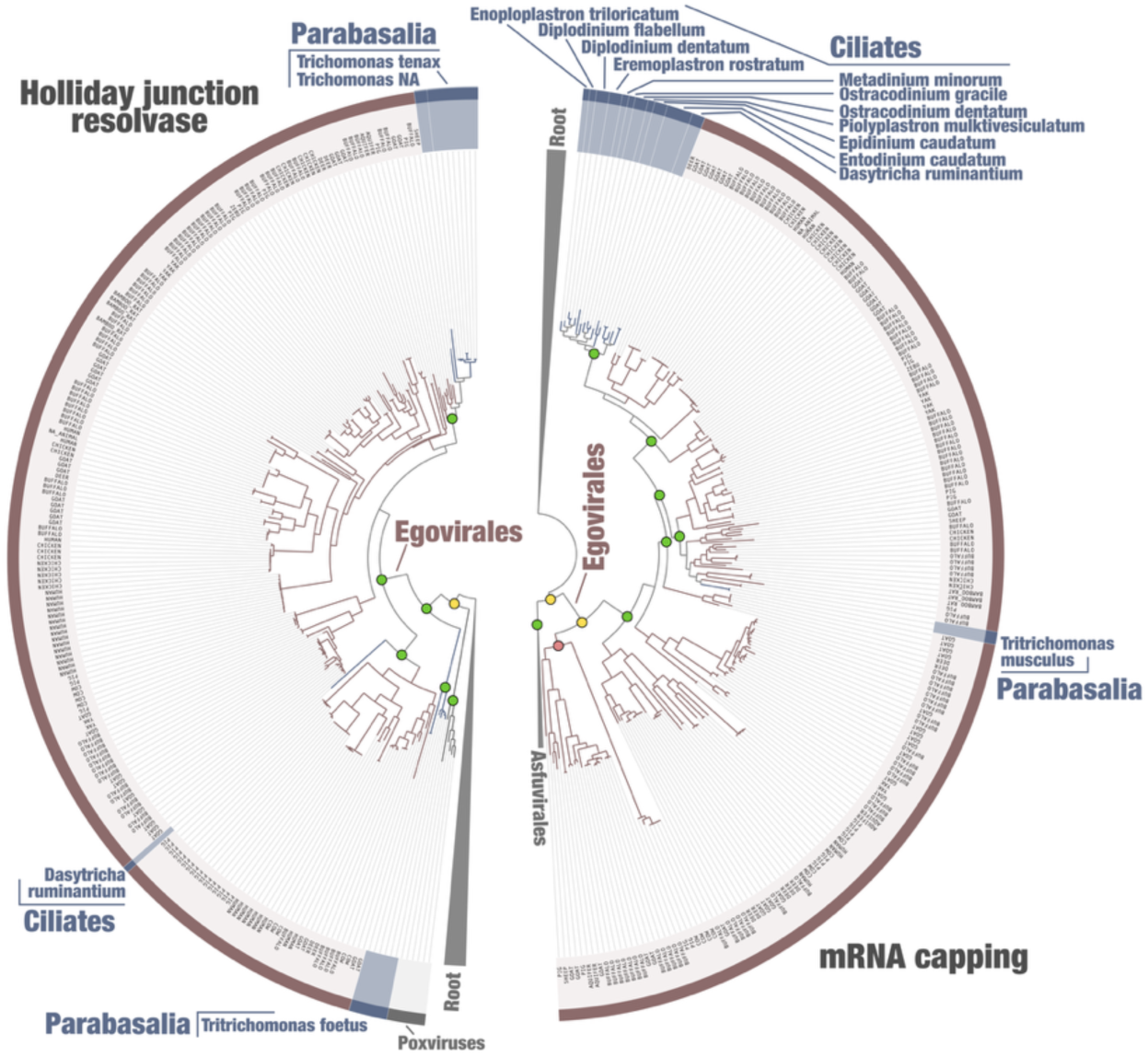
Phylogenies of the Holliday Junction resolvase and mRNA capping enzyme. Left tree corresponds to the maximum-likelihood phylogeny of the Holliday Junction of *Nucleocytoviricota* and eukaryotes (1,050 taxa; 277 sites; model Q.YEAST+F+R9), rooted at the level of *Egovirales*. Right tree corresponds to the maximum-likelihood phylogeny of the mRNA capping of *Nucleocytoviricota* and eukaryotes (607 taxa; 807 sites; model Q.PFAM+F+R9), rooted at the level of *Egovirales*. Green dots at nodes in the tree highlight high supports (approximate likelihood ratio (aLRT) ≥ 80 and ultrafast bootstrap approximation (UFBoot) ≥ 95; see Methods). Yellow dots represent moderate supports (aLRT >= 80 or UFBoot >= 95).

**Figure S8:**
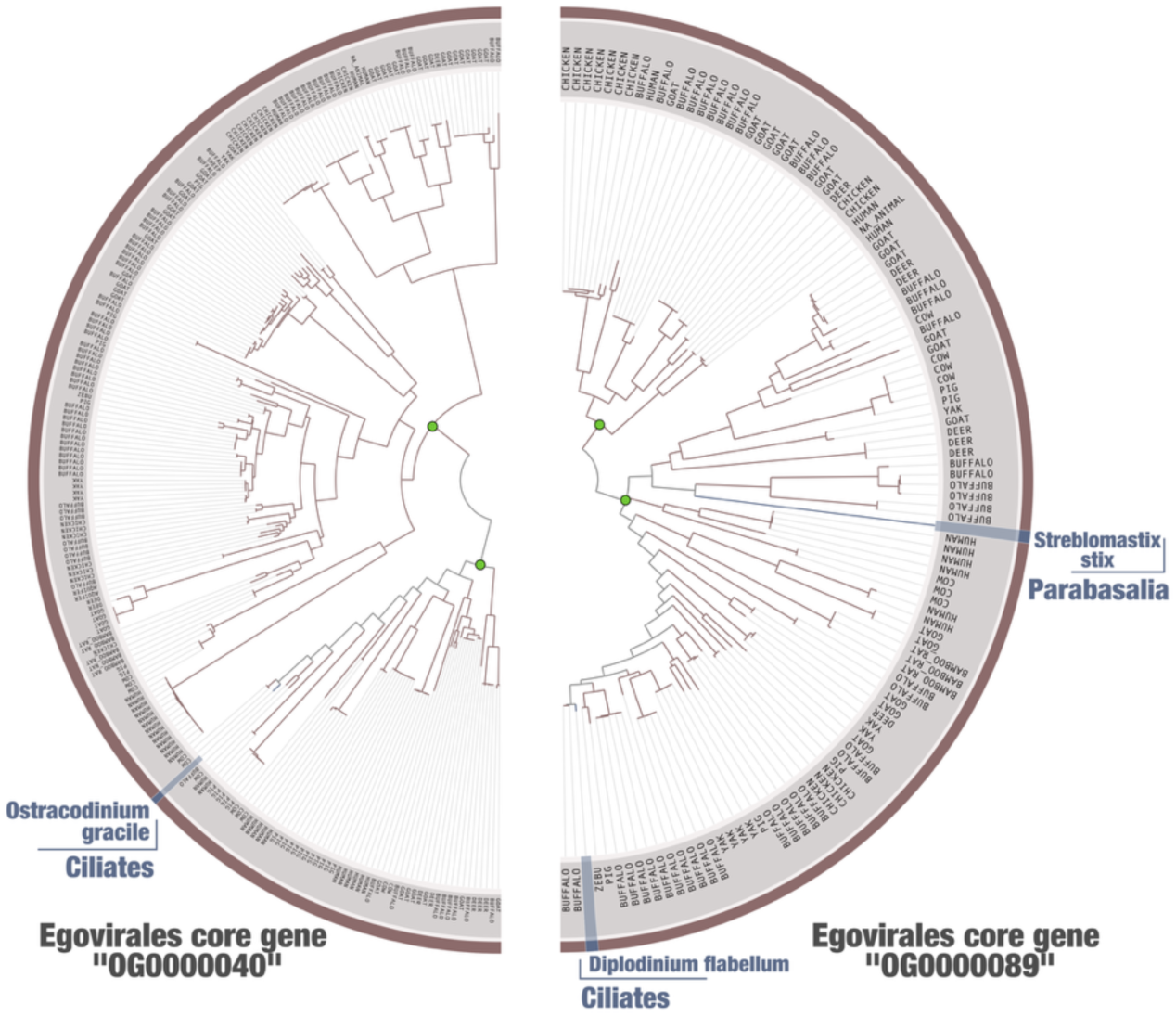
Phylogenies of two egovirus core genes of unknown function. Left tree corresponds to the maximum-likelihood phylogeny of egovirus core gene OG0000040 (219 taxa; 398 sites; model Q.INSECT+F+R5). Right tree corresponds to the maximum-likelihood phylogeny of egovirus core gene OG0000089 (116 taxa; 383 sites; model Q.PFAM+F+R4). Green dots at nodes in the tree highlight high supports (approximate likelihood ratio (aLRT) ≥ 80 and ultrafast bootstrap approximation (UFBoot) ≥ 95; see Methods).

