## Supplementary Information for "Closest relatives of poxviruses replicate in the digestive system of humans and animals worldwide"

### Supplemental Information

#### 1- Recovery of egovirus environmental genomes at scale

The identification of a deep-branching clade of RNApolA from the mOTUs[1] database was the starting point of this environmental genomic survey. This deep-branching clade allowed the characterization of a first set of RNApolA-positive Egovirales contigs, 12 of which turned out to be complete non-redundant genomes (average nucleotide identity <98%) based on the occurrence of 55 bp inverted terminal repeats. These complete genomes occur in the digestive system of humans, buffalos, goats, chicken, deer, and pigs. Using Orthofinder[2], we identified 58 core genes present in at least 10 of them. These core genes were functionally annotated using HHpred[3] profile-profile comparisons and ColabFold[4] (3D structure predictions and comparisons). In the case of the DJR-MCP of *Egovirales*, ColabFold[4] successfully predicted its 3D structure and confirmed the presence of the double-jelly roll (DJR) protein components characteristic of the phylum *Nucleocytoviricota*.

Among the 58 core genes of *Egovirales*, 51 are single-copy core genes relevant to assessing the completion and redundancy of environmental genomes. Using this *Egovirales*-centric collection of single copy core genes, the 12 complete non-redundant genomes are estimated to be on average 96% complete and 1% redundant. Among the same 58 core genes of *Egovirales*, 42 lack signal in other lineages of nucleocytoviruses (still based on Orthofinder) and were labelled as *Egovirales*-specific core genes. These genes are relevant to automatically identify Egovirales contigs from metagenomic assemblies. Tables S3 and S4 provides information on the different set of *Egovirales* core genes.

Among the mOTUs[1] automatically generated bins with signal for the RNApolA or DJR-MCP of *Egovirales*, we used 42 *Egovirales*-specific core genes (those rarely occur in other lineages of nucleocytoviruses) to confidently extract Egovirales contigs and create environmental genomes. Simply, contigs lacking signal for any of these 42 *Egovirales*-specific core genes were considered likely contaminants and removed. We subsequently used the 51 *Egovirales* single-copy core genes to assess their completion and redundancy. Environmental genomes with low quality metric (completion minus redundancy) were excluded. This method allowed us to effectively characterize *Egovirales* environmental genomes at scale from the mOTUs database (tens of thousands of metagenomic assemblies from a broad range of biomes) and to estimate their quality scores despite the lack of any reference genome from culture for this newly identified viral order. Clearly, the early identification of 12 complete non-redundant genomes was instrumental for the success of this environmental genomic survey.

#### 2- The virion morphogenesis module of egoviruses

Analysis of the virion morphogenesis module of egoviruses sheds light on the key features of the egovirus virions. Egoviruses encode the major components of the virion morphogenetic module conserved across *Nucleocytoviricota*, including poxviruses and asfuviruses (Figure 1). The core components of this module are the double jelly-roll (DJR) major capsid protein[5] (MCP; Figure 2), the A32-like genome packaging ATPase, and Ulp1-

like capsid maturation protease. The DJR-MCP characteristic of *Nucleocytoviricota* was particularly challenging to identify in *Egovirales* due to its extensive protein sequence divergence compared to the closest reference proteins. Our initial searches using a dedicated HMM, which successfully recovered DJR-MCPs from all previously described families of the phylum *Nucleocytoviricota*[6,7], systematically failed to recognize this protein in egoviruses. Eventually, HHpred[3] profile-profile comparisons yielded a partial match to the DJR-MCP of African swine fever virus, and structural modeling with ColabFold[4] confirmed that the predicted egovirus MCP has the DJR fold. The *Egovirales* DJR-MCPs are unusually large (average of 806 amino acids among the complete *Egovirales* genomes) and the region facing the exterior of the capsid is highly diversified among egoviruses. These trends echo the previous observations on the large MCPs of faustoviruses from the order *Asfuvirales*[8]. However, *Egovirales* and *Chitovirales* are firmly established as sister clades in the DJR-MCP phylogeny, despite the substantial protein length differences between the two orders. In addition, egoviruses encode homologs of the major virion proteins exclusive to asfuviruses, namely, structural polyproteins pp220 and pp62. In African swine fever virus, both polyproteins are processed into multiple structural proteins, which play important roles in the formation of multilayered capsids. One of the major proteolytic products of pp220, p150, that is conserved in egoviruses, has an α-helical fold and forms an internal icosahedral shell located inside of a larger, external icosahedral capsid formed from the DJR-MCP[9]. So far, the extent of protein sequence diversification among egoviruses prevented the prediction of the high-confidence 3D structure for the p150 homolog. Structure-based searches allowed identification of the homologs of asfuviral core shell proteins p15 and p35, two of the three proteolytic products of pp62 polyprotein. The p15 and p35 bridge the viral nucleoprotein complex and the internal capsid to the lipid membrane sandwiched between the inner and outer icosahedral shells[10,11]. Collectively, the presence of these three structural proteins suggests that egoviruses form icosahedral multilayered capsids that are similar to those of *Asfuvirales*[9], but differ from the brick-shaped capsids of poxviruses[12].

#### 3- High prevalence of egoviruses in buffalos

Buffalos had by far the highest *Egovirales* positivity rate among the surveyed vertebrate species, with viruses covering a wide range of distantly related clades (Figure 2 of the main text). The considered cohorts cover various Chinese provinces and include three buffalo breeds and different segments of their digestive system[13], allowing for a more refined survey of the occurrence of egoviruses within the gut microbiome of an herbivorous animal species (Table S6). First, we found up to four distinct egoviruses co-occurring in the same sample, contrasting with the human gut microbiome, where no more than one egovirus was detected in the same individual. Second, the four ruminant stomach compartments displayed the highest prevalence of egoviruses: 79% of positive samples for the reticulum, 45% for the abomasum, 38% for the omasum, and 20% for the rumen (Supp. Information Table 1). Egoviruses were also detected in the cecum (n=19%) and distal colon (n=10%). Finally, the positivity rate reached 25.8% among fecal samples. Based on the fecal samples, egoviruses were detected in all seven Chinese provinces with a positivity rate ranging from 6.7% to 52%, demonstrating the widespread occurrence of *Egovirales* in the buffalo gut microbiome.

| | Nb of samples | Samples with viral detection | Positivity rate |
|---|---|---|---|
| Reticulum | 28 | 22 | 79% |
| Abomasum | 22 | 10 | 45% |
| Omasum | 32 | 12 | 38% |
| Feces | 399 | 103 | 26% |
| Rumen | 129 | 26 | 20% |
| Cecum | 31 | 6 | 19% |
| Colon | 30 | 3 | 10% |
| Jejunum | 24 | 0 | 0% |
| Total | 695 | 179 | 26% |

**Egovirus occurence inside the Buffalo**

| | Nb of samples | Samples with viral detection | Positivity rate |
|---|---|---|---|
| China - Hainan | 31 | 16 | 52% |
| Myanmar | 31 | 13 | 42% |
| China - Yunnan | 62 | 25 | 40% |
| China - Anhui | 44 | 11 | 25% |
| China - Guangxi | 175 | 34 | 19% |
| China - Henan | 41 | 3 | 7% |
| China - Hubei | 15 | 1 | 7% |

**Egovirus occurence across regions**

**Supp. information Table 1: Occurrence of egoviruses in Buffalos.**

#### 4- Gene flux between egoviruses and unicellular eukaryotes

Our overall analysis on gene flux between egoviruses and eukaryotes relies on complementary approaches.

- We functionally annotated (Pfam) and ran Orthofinder on predicted proteins from all the egovirus environmental genomes.
- We compared each egovirus protein to a large protein database that covers the three domains of life plus the viruses and removed all eukaryotic hits for which a better hit (e.g., for a Bacteria) occurred. This allowed us to focus on a clean egovirus-eukaryotic signal, which depicted a strong connection between egoviruses and two distantly related unicellular eukaryotic clades: *Parabasalia* (Metamonada) and *Trichostomatia* (Ciliates). As a critical perspective, none of the other lineages of nucleocytoviruses display such a link to these two eukaryotic lineages (Supp. Information Table 2).
- We explored the functions and protein clusters of egoviruses associated with this gene flux (Egovirus-*Parabasalia* and Egovirus-*Trichostomatia*).
- We performed a complementary analysis, in which proteins of Metamonada and Ciliates are compared to that of nucleocytoviruses (including the egoviruses). This work suggests that egovirus hits are spread in the eukaryotic chromosomes, and revealed that one of the core gene of egoviruses (KilA) is associated with polintons spread in the genomes of *Parabasalia* and especially *Trichomonas vaginalis*.
- We performed phylogenies for the most prominent functions connecting egoviruses and the two eukaryotic clades (Figures S6, S7 and S*), as well as a phylogeny of KilA to connect egoviruses and polintons (Figure 3).

| | Ego | Chito | Asfu | Alga | Imiter | Pandora | Pimasco | Mirus |
|---|---|---|---|---|---|---|---|---|
| Nb of genomes | 224 | 43 | 39 | 324 | 1339 | 121 | 212 | 111 |
| Nb of genes | 43500 | 8367 | 12048 | 84107 | 540524 | 55750 | 54473 | 22825 |
| Euk best hits | 2417 | 712 | 1012 | 5335 | 22252 | 2897 | 2455 | 896 |
| Amoebozoa | 6.3 | 0.6 | 4.4 | 2.6 | 5.9 | 12.9 | 11.6 | 2.8 |
| Viridaplantae | 1.2 | 0.3 | 8.6 | 62.9 | 16.4 | 11.1 | 8 | 11.5 |
| Metamonada | 42.4 | 8 | 4.1 | 2.2 | 5.4 | 4.2 | 3.7 | 8.7 |
| Parabasalia | 37.4 | 6.2 | 0.8 | 0.5 | 1.7 | 0.7 | 0.9 | 3 |
| Opistokonta | 6 | 89.3 | 34.6 | 19.1 | 30.9 | 16.5 | 38.5 | 28.5 |
| Stramenopiles | 1.1 | 0.3 | 33.6 | 5.1 | 16.3 | 23.4 | 16.3 | 25.4 |
| Rhizaria | 0 | 0 | 0.7 | 0.3 | 1.5 | 0.6 | 1.6 | 1.6 |
| Ciliophora | 42.3 | 1 | 5.8 | 2.6 | 9.2 | 4.2 | 10.5 | 5.7 |
| Trichostomatia | 42.2 | 0.6 | 3.6 | 2 | 6.7 | 2.5 | 8.7 | 4.9 |
| Rhodophyta | 0.1 | 0 | 0.6 | 0.5 | 1.1 | 0.8 | 1.1 | 1.7 |
| Haptista | 0 | 0 | 1.2 | 1.3 | 2.5 | 20.8 | 1.6 | 3.3 |
| Cryptista | 0 | 0 | 0.8 | 0.7 | 3.1 | 0.3 | 0.9 | 1 |
| Discobia | 0.3 | 0.1 | 2.5 | 0.6 | 2.6 | 0.8 | 2.3 | 1.1 |
| Other eukaryotes | 0 | 0 | 3.2 | 2.2 | 5.2 | 4.4 | 3.8 | 8.7 |

Percent hits among eukaryotes

**Supp. information Table 2: Gene flux between lineages of nucleocytoviruses (plus the phylum Mirusviricota) and major lineages of eukaryotes.** The table shows a spike of signal connecting egoviruses with both *Parabasalia* and *Trichostomatia*.
